# Draft Genome Sequence of *Candida auris* Strain LOM, a Human Clinical Isolate from Greater Metropolitan Houston, Texas

**DOI:** 10.1101/629741

**Authors:** S. Wesley Long, Randall J. Olsen, Hoang A. T. Nguyen, Matthew Ojeda Saavedra, James M. Musser

## Abstract

*Candida auris* is an emerging pathogen with considerable public health importance. We present the draft genome sequence of a strain recently cultured from the urine of a patient hospitalized in the greater Houston metropolitan region. Two combined Oxford Nanopore sequencing runs provided sufficient data to rapidly generate a draft genome.

## ANNOUNCEMENT

*Candida auris* recently has emerged worldwide and caused outbreaks in healthcare facilities (1, 2). An isolate of *C. auris* cultured from the urine of a patient from the Houston metropolitan region was identified by Brucker MALDI-TOF analysis using the RUO database. Given the potential infection control and public health importance of this isolate, we rapidly characterized its genome using two Oxford Nanopore GridION runs. Initial DNA extraction was performed using ballistic lysis with FastPrep matrix B beads and the Qiagen DNA Blood/Tissue Kit. The first GridION run was performed with the Rapid Barcoding Kit from Oxford Nanopore (SQK-RBK004). The second run was performed using DNA extracted using FastPrep matrix Y beads, the Qiagen DNA Blood/Tissue Kit. The Ligation Sequencing Kit from Oxford Nanopore (SQK-LSK109) with a long read wash step to select for longer reads as suggested in the manufacturer’s protocol. Initial genome assembly of GridION data from the first 16 hours of the first sequencing run using Unicycler v0.4.3 yielded 11 contigs in the aggregate corresponding to 12,237,085 bps (3). After completion of the second GridION run, the long reads from both runs were combined and filtered using FiltLong v0.1.1 (https://github.com/rrwick/Filtlong) with a minimum read length cutoff at 5 kb and a target of 100x-fold coverage. Unicycler v0.4.3 was used to assemble the genome from the long reads. The final assembly had 9 contigs with a total length of 12,293,266 bps. The largest contig is 4,297,164 bps, and the N50 is 2,304,466 bps. The average GC content is 45.19%. This work was approved by our Institutional Review Board (IRB1010-0199).

We compared our assemblies with reference genome assemblies deposited in GenBank using progressiveMauve v2.4.0 and discovered that our 7 longest contigs correspond to the expected 7 chromosomes present in other *C. auris* strains (4). Phylogenetic analysis using mummer v4.0 to compare against the available reference genomes shows that *C. auris* strain LOM is most closely related to strain B11221, a representative strain from clade III, a lineage with many South African strains (5, 6). This relationship was confirmed using prephix and phrecon to generate an alignment for FastTree to create a Neighbor Joining phylogenetic tree (https://github.com/codinghedgehog/). The *C. auris* strain LOM genome has the *ERG11* gene with a F126T amino acid replacement that is common to clade III strains. (6).

Although an in-depth epidemiological investigation had not been completed at the time of manuscript submission, the patient had not recently traveled outside the United States, had a history of long-term care at outside facilities, and had very recently been transferred to hospital A from another facility. The organism was classified as present on admission. This work further demonstrates the usefulness of real-time long read whole genome sequencing to rapidly provide relevant information concerning emerging pathogens of significant concern (7, 8). The availability of this high-quality draft genome sequence serves as a useful resource if additional isolates are recovered in greater Houston, a metropolitan area with an international, ethnically diverse population of approximately 6 million. The sequence data also will assist translational research efforts designed to more fully understand the molecular mechanisms underlying host interactions in an emerging pathogen that has substantial detrimental public health potential.

## Data availability and accession numbers

The BioProject accession for the draft genome sequence data of *C. auris* strain LOM is <u>PRJNA540998</u>. The GridION runs are present in the SRA (<u>SRR9017243</u> and <u>SRR9017244</u>). The assembled draft genome can be found in GenBank (<u>SZYF00000000</u>).

## ACKNOWLEDGMENTS

We thank the technical staff in the Houston Methodist Hospital Clinical Microbiology Laboratory for assistance. We thank Andrew Pann for his development and continued support of the prephix and phrecon tools. Supported by funds from the L. E. Simmons Family Foundation and Fondren Foundation to J.M.M.

